# Methylome-dependent transformation of *emm*1 group A streptococci

**DOI:** 10.1101/2023.04.10.536284

**Authors:** Joana Alves, Joshua D Rand, Alix BE Johnston, Connor Bowen, Nicola N Lynskey

## Abstract

Genetic intractability presents a fundamental barrier to the manipulation of bacteria, hindering advancements in microbiological research. Group A *Streptococcus* (GAS), a lethal human pathogen currently associated with an unprecedented surge of infections world-wide, exhibits poor genetic tractability attributed to the activity of a conserved type 1 restriction modification system (RMS). RMS detect and cleave specific target sequences in foreign DNA that are protected in host DNA by sequence-specific methylation. Overcoming this “restriction barrier” thus presents a major technical challenge. Here we demonstrate for the first time that different RMS variants expressed by GAS give rise to genotype-specific and methylome-dependent variation in transformation efficiency. Further, we show that the magnitude of impact of methylation on transformation efficiency elicited by RMS variant TRD_AG_, encoded by all sequenced strains of the dominant and upsurge-associated *emm*1 genotype, is 100-fold greater than for all other TRD tested and is responsible for the poor transformation efficiency associated with this lineage. In dissecting the underlying mechanism, we developed an improved GAS transformation protocol whereby the restriction barrier is overcome by addition of the phage anti-restriction protein Ocr. This protocol is highly effective for TRD_AG_ strains including clinical isolates representing all *emm*1 lineages, and will expedite critical research interrogating the genetics of *emm*1 GAS, negating the need to work in an RMS-negative background. These findings provide a striking example of the impact of RMS target sequence variation on bacterial transformation, and the importance of defining lineage-specific mechanisms of genetic recalcitrance.

**Importance:** Understanding the mechanisms by which bacterial pathogens are able to cause disease is essential to enable the targeted development of novel therapeutics. A key experimental approach to facilitate this research is the generation of bacterial mutants, through either specific gene deletions or sequence manipulation. This process relies on the ability to transform bacteria with exogenous DNA designed to generate the desired sequence changes. Bacteria have naturally developed protective mechanisms to detect and destroy invading DNA, and these systems severely impede the genetic manipulation of many important pathogens, including the lethal human pathogen group A *Streptococcus* (GAS). Many GAS lineages exist, of which *emm*1 is dominant among clinical isolates. Based on new experimental evidence, we identify the mechanism by which transformation is impaired in the *emm*1 lineage and establish an improved and highly efficient transformation protocol to expedite the generation of mutants.

## Observation

Genetic manipulation of pathogenic bacteria is an essential tool to characterise virulence mechanisms and develop new therapies. However, research into certain pathogens is hampered by their inherent resistance to genetic transformation. One such species is GAS, an obligate human pathobiont and the causative agent of a diverse array of infections ranging from pharyngitis and scarlet fever to necrotising fasciitis^1^. Genotype *emm*1 GAS are responsible for an ongoing and unprecedented global surge in infections, the molecular basis for which remains unknown^2–5^. The development of an improved transformation protocol for *emm*1 GAS is thus essential to expedite critical research into the pathophysiology of this genotype.

The genetic recalcitrance of GAS has been linked to the activity of a chromosomally-encoded type 1 restriction modification system (RMS) that is conserved across all genotypes^6–8^. Type 1 RMS comprise three genes, a DNA restriction endonuclease (*hsdR*), methyl-transferase (*hsdM*) and DNA specificity protein (*hsdS*) which combined form a holoenzyme with methyl-transferase and DNA-cleaving activity^9^. The HsdS protein defines the target DNA sequence motif through two distinct 5’ and 3’ target recognition domains (TRD). Thirteen distinct GAS TRD combinations have been identified, each of which targets a unique DNA motif and is associated with a specific subset of genotypes^6^. Interestingly, all *emm*1 strains are associated with a single TRD combination, designated TRD_AG_^6^. While targeted deletion of the RMS and naturally-occurring, inactivating mutations have been demonstrated to enhance transformation efficiency of three GAS genotypes^6–8^, a direct comparison of the impact of the different methylation patterns conferred by each TRD combination on transformation efficiency has not been performed.

### Methylation-dependent DNA restriction drives the low transformation efficiency of *emm*1 GAS

In order to determine whether GAS transformation efficiencies differ due to the activity of different TRD variants, transformation efficiencies were quantified for strains representing genotypes most frequently subjected to genetic manipulation for six TRD combinations (Figure 1A, Table S1). Interestingly, the tested genotypes segregated into two discrete groups with low (10^4^ cfu/μg DNA, *emm*4/TRD_AF_, *emm*1/TRD_AG_, *emm*5/TRD_FA_) or high (10^6^ cfu/μg DNA, *emm*89/TRD_BG_, *emm*49/TRD_CF_, *emm*18/TRD_DA_) transformation efficiencies (Figure 1B). We hypothesised that this difference was a methylation-dependent phenomenon driven by variation in the target sequence for each TRD combination.

**Figure 1:**
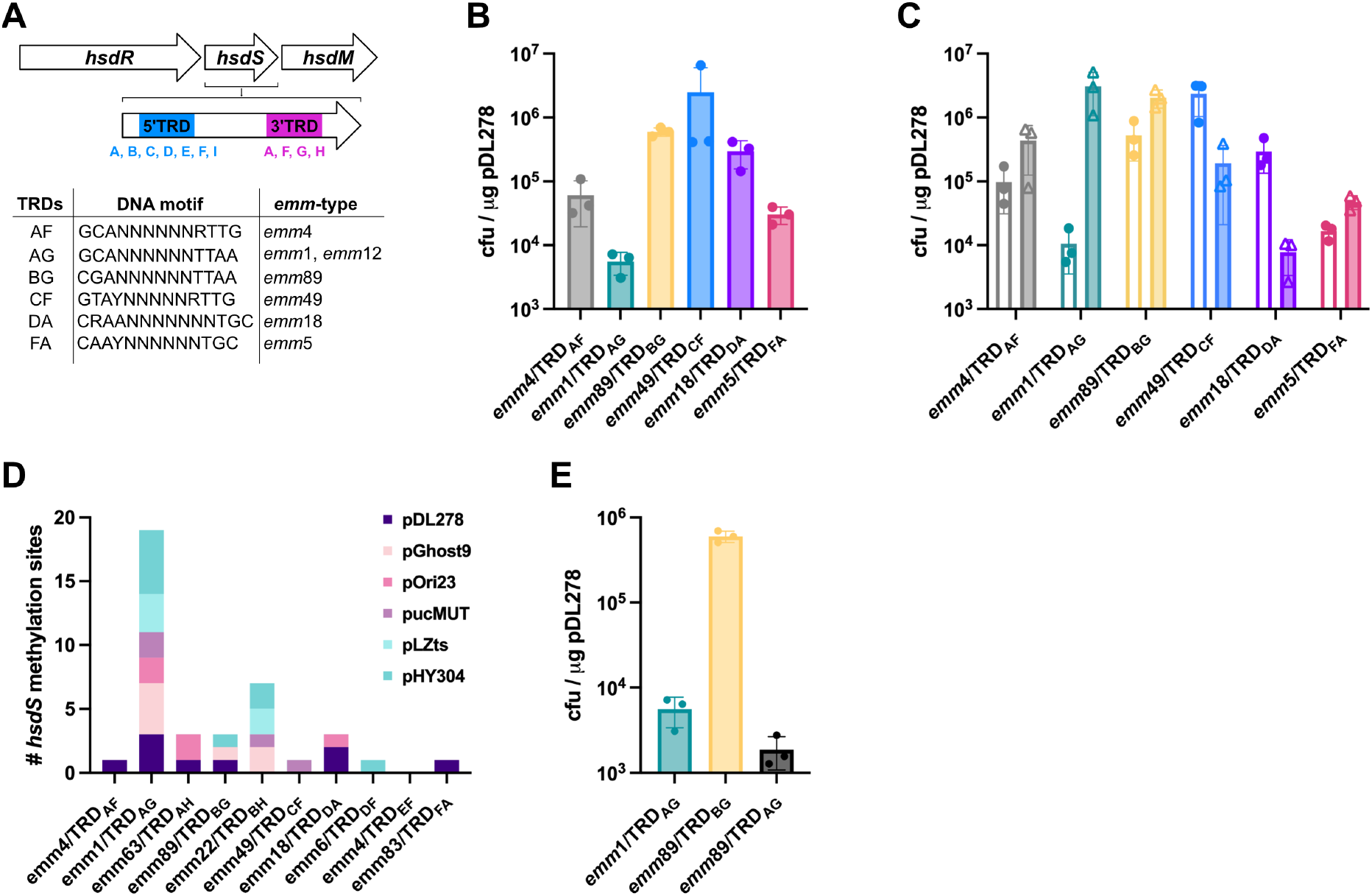
DNA methylation defines the reduced transformation efficiency of *emm*1 GAS. (**A**) Schematic representation of the GAS type 1 RMS and location of TRDs (5’ = blue, 3’ = pink) in the *hsdS* gene (M5005_Spy1622). Each TRD variant identified to date listed underneath. Table shows six TRD combinations associated with GAS genotypes most frequently subjected to experimental genetic manipulation with details of DNA target motifs^6^ and representative associated genotypes used in this study. (**B**) Transformation efficiency of a representative isolate from each of the six TRD combinations highlighted in (A) with plasmid pDL278. Transformation efficiency of three TRD combinations (*emm*4/TRD_AF_, *emm*1/TRD_AG_, *emm*5/TRD_FA_) is reduced compared with the other strains tested (*emm*89/TRD_BG_, *emm*49/TRD_CF_, *emm*18/TRD_DA_). (**C**) Comparison of transformation efficiency of a representative isolate from each of the 6 TRD combinations highlighted in (A) with DH5α-(clear bars) or self-methylated (filled bars) plasmid pDL278. Transformation efficiency of *emm*1/TRD_AG_ isolate was enhanced with self-methylated plasmid. (**D**) Quantification of DNA target motifs for all GAS TRDs in each of six commonly used plasmids for genetic manipulation of streptococcal pathogens. The target sequence for *emm*1/TRD_AG_ (5’-GCANNNNNNTTAA-3’) was identified more frequently than all other target sequences across all six plasmids. (**E**): Comparison of transformation efficiency of *emm*1/TRD_AG_ and *emm*89/TRD_BG_ with the “TRD-swap” strain, *emm*89/TRD_AG_ with plasmid pDL278 purified from DH5α. Transformation efficiency of the *emm*89/TRD_BG_ strain was reduced to levels equivalent to *emm*1/TRD_AG_ following the swap of TRD_B_ to TRD_A_.

All transformations were performed using plasmid DNA purified from the DH5α strain of *E. coli* lineage K12. K12 *E. coli* encode three methyl-transferases (Dam_G<u>A</u>TC, Dcm_C<u>C</u>WGG and EcoKI_AACNNNNNNG<u>T</u>GC/GC<u>A</u>CNNNNNNGTT) that target DNA sequences distinct from all known GAS TRDs. Plasmid purified from DH5α will thus be susceptible to cleavage by all GAS TRD variants, where target sites are present. In order to ascertain whether TRD-specific methylation was responsible for the observed differences in transformation efficiency between GAS TRD variants, we tested whether self-methylated plasmid purified from each GAS strain, and thus protected from self-RMS cleavage, impacted DNA uptake (Figure 1C). Strikingly, *emm*1/TRD_AG_ transformation efficiency was increased 100-fold using self-methylated plasmid. Surprisingly, the difference in transformation efficiency between self and DH5α originated plasmid was less pronounced or absent for all other strains tested, including those with an equivalent transformation efficiency to that of *emm*1/TRD_AG_ using DH5α-purified plasmid. This led us to hypothesise that the activity of the type 1 RMS is higher for *emm*1/TRD_AG_ strains or that more TRD_AG_ recognition motifs are present in the plasmid DNA.

### Unique methylation motif of TRD_AG_ is common in plasmids used for bacterial genetic engineering

The absence of restriction targets in plasmid DNA sequences is an important RMS evasion strategy^10^. In order to determine whether the absence of target sequences could explain the unique methylation-associated phenotype for *emm*1/TRD_AG_ GAS, we first compared the frequency of each known GAS TRD recognition motif in plasmid pDL278^11^ used for all experiments, and then expanded our analysis to include five additional commonly used laboratory plasmids^12–16^ (Table S2). Unexpectedly, the *emm*1/TRD_AG_ recognition sequence was overrepresented across all six plasmids (Figure 1D), and thus likely contributes to the low transformation efficiency observed for *emm*1/TRD_AG_ GAS.

While over-representation of the *emm*1/TRD_AG_ recognition site may completely explain the low transformation efficiency of this lineage, the low efficiency observed for *emm*4/TRD_AF_ and *emm*5/TRD_FA_ strains indicates that other factors also contribute to genetic recalcitrance. We hypothesised that lineage-specific variation in RMS gene sequences or expression levels may also contribute to resistance to transformation, and went on to generate a “TRD-swap” strain in our representative *emm*89/TRD_BG_ isolate to quantitatively assess this possibility. The *emm*1/TRD_AG_ and *emm*89/TRD_BG_ *hsdS* alleles share the same 3’ TRD. Allelic exchange mutagenesis was performed to swap the *emm*89 5’ TRD_B_ with TRD_A_ to generate the TRD_AG_ allele in the *emm*89/TRD_BG_ background, giving rise to strain *emm*89/TRD_AG_. This swap reduced the transformation efficiency of the *emm*89 strain by 100-fold to levels observed for *emm*1/TRD_AG_ (Figure 1E), a result that strongly implicates over-representation of the *emm*1/TRD_AG_ motif as the basis for poor transformation efficiency of *emm*1/TRD_AG_ strains.

### Inhibition of *emm*1/TRD_AG_ type 1 RMS with phage protein Ocr enhances transformation efficiency

Having shown that the reduced transformation efficiency associated with *emm*1/TRD_AG_ could be explained by over-representation of TRD_AG_ target motifs in commonly used plasmids, we went on to determine whether inhibition of the type 1 RMS with the phage anti-restriction protein Ocr^17^ could improve transformation efficiency of this genotype without the need to generate an isogenic RMS deletion mutant. Addition of Ocr protein (50 ng/μl) to the electroporation reaction enhanced the efficiency 100-fold following transformation with DH5α-purified plasmid, equivalent to the highest efficiencies observed for *emm*89/TRD_BG_ strains but had no effect on transformation with self-methylated plasmid (Figure 2A).

**Figure 2:**
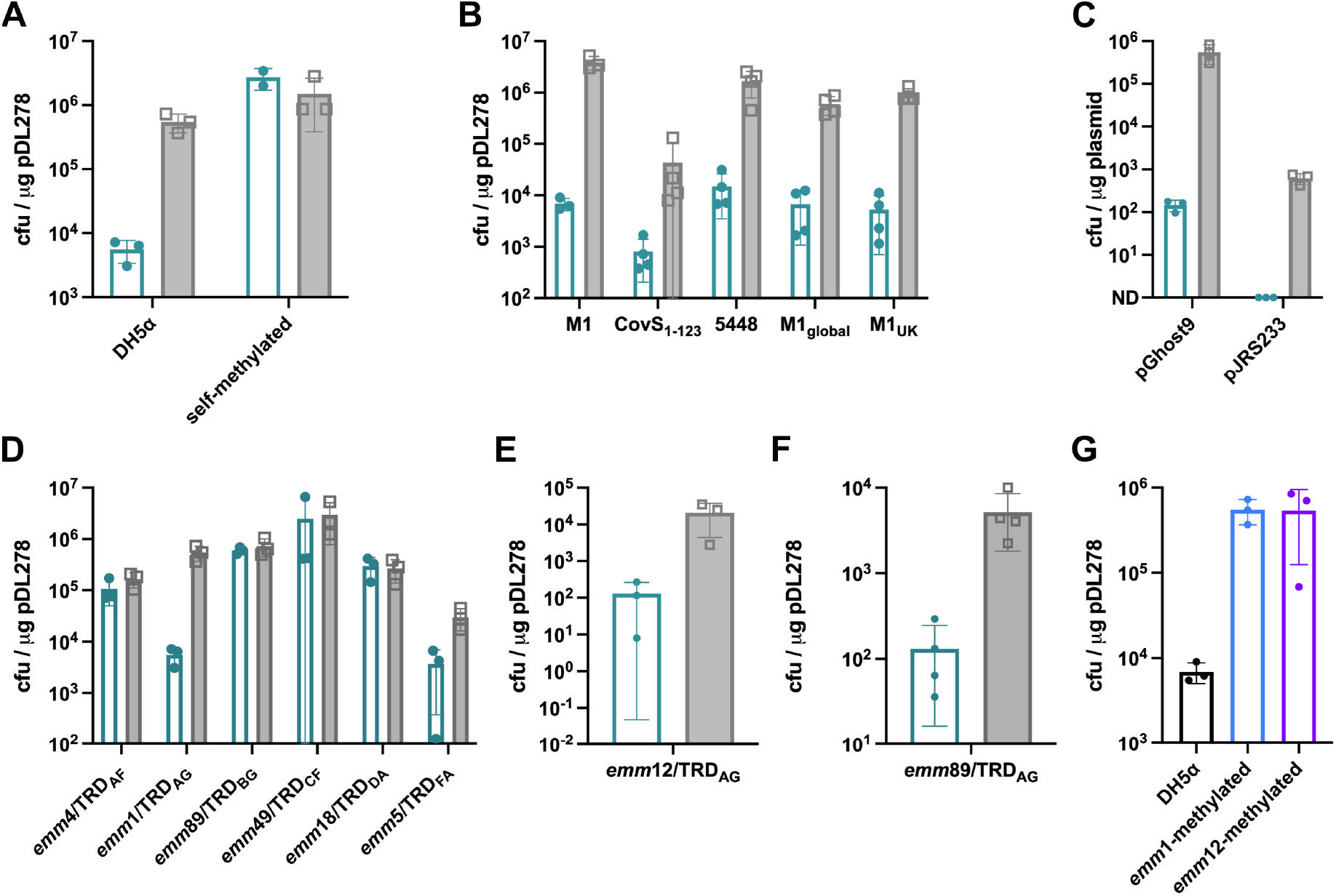
Transformation efficiency of *emm*1 GAS is enhanced 100-fold by inhibition of HsdR with phage protein Ocr. (**A**) Quantification of transformation efficiency of *emm*1/TRD_AG_ with DH5α- or self-methylated plasmid (pDL278) in the presence/absence of recombinant Ocr protein (Clear green bars = - Ocr; filled grey bars = +Ocr). Ocr enhanced the transformation efficiency of *emm*1/TRD_AG_ with DH5α-methylated pDL278 to levels equivalent to self-methylated plasmid, however had no impact on efficiency of transformation with self-methylated plasmid. (**B**) Quantification of transformation efficiency of 5 *emm*1 isolates representing all major clinically-relevant and globally-disseminated strains/lineages with DH5α-purified plasmid pDL278 +/- recombinant Ocr (Clear green bars = - Ocr; filled grey bars = +Ocr). Ocr improved transformation efficiency with plasmid pDL278 100-fold for all strains. (**C**) Quantification of transformation efficiency of strain *emm*1/TRD_AG_ with chromosomal allelic exchange vectors pGhost9 and pJRS233. Plasmids were purified from DH5α and transformed +/- recombinant Ocr protein(Clear green bars = - Ocr; filled grey bars = +Ocr). Addition of Ocr enhanced transformation efficiency 10^4^- and 10^3^-fold respectively. (**D**) Quantification of transformation efficiency of a representative isolate from each of the six TRD combinations highlighted in Figure 1A with plasmid pDL278 purified from DH5α +/- recombinant Ocr protein (Clear green bars = - Ocr; filled grey bars = +Ocr). Addition of Ocr had no effect on transformation efficiency any strain other than *emm*1/TRD_AG_. (**E**) Quantification of transformation efficiency of *emm*12/TRD_AG_ with plasmid pDL278 purified from DH5α +/- recombinant Ocr (Clear green bars = - Ocr; filled grey bars = +Ocr). Ocr enhanced transformation efficiency 100-fold. (**F**) Quantification of transformation efficiency of *emm*89/TRD_AG_ with plasmid pDL278 purified from DH5α +/- recombinant Ocr (Clear green bars = - Ocr; filled grey bars = +Ocr). Ocr enhanced transformation efficiency 100-fold. (**G**) Quantification of transformation efficiency of *emm*1/TRD_AG_ with plasmid pDL278 purified from DH5α, *emm*1/TRD_AG_ and *emm*12/TRD_AG_. Transformation of *emm*1/TRD_AG_ with plasmid purified from either GAS strain expressing TRD_AG_ was enhanced by 100-fold.

In order to confirm that this was not a strain-specific phenomenon, we expanded the isolates tested to include *emm*1 strains representing the most common genetic variants; a naturally occurring CovS_1_-123 mutant^18^, M1T1 strain 5448 (USA origin)^19^ and strains representing the two dominant lineages circulating currently and responsible for the current upsurge in GAS infections globally, M1_global_^5^ and M1_UK_^5^. Transformation of all *emm*1 strains, each encoding TRD_AG_, was improved by the same magnitude following incorporation of Ocr into the transformation reaction (Figure 2B). This result demonstrates that the transformation efficiency of multiple *emm*1 clinical isolates from diverse lineages is greatly enhanced using this protocol.

In order to confirm that this effect was not specific to the pDL278 plasmid, we performed similar experiments with plasmids pGhost9^12^ and pJRS233^20^, frequently used as suicide vectors. Addition of Ocr protein enhanced the transformation efficiency of *emm*1/TRD_AG_ with both plasmids by an even greater magnitude than that observed for pDL278 (Figure 2C).

### Ocr enhancement of GAS transformation is restricted to GAS expressing TRD_AG_

We went on to ascertain whether the addition of Ocr was sufficient to enhance the transformation efficiency of strains representing other TRD combinations. As observed for transformation with self-methylated plasmid (Figure 1B), Ocr had only a marginal impact on transformation efficiency of most non-*emm*1/TRD_AG_ genotypes (Figure 2D). Importantly, similar to *emm*1 GAS, the efficiency of DNA uptake by an *emm*12/TRD_AG_ strain (Figure 2E) and the *emm*89/TRD_AG_ (Figure 2F) were enhanced 2 log-fold following addition of Ocr protein, indicating that this phenotype is methylation-dependent. This conclusion is further supported by the observation that transformation of *emm*1/TRD_AG_ GAS was similarly enhanced using plasmid purified from and methylated by either *emm*1/TRD_AG_ or *emm*12/TRD_AG_ GAS (Figure 2G).

Together these data demonstrate that the type 1 RMS prevents efficient transformation of *emm*1 GAS due to over-representation of the target methylation site in plasmids, and that this restriction barrier can be overcome by addition of the phage protein Ocr. We go on to establish an experimental protocol for enhanced transformation of TRD_AG_ GAS that is effective for diverse clinical *emm*1 strains.

## Data availability

All relevant data are within the manuscript and Supporting Information files.

## Acknowledgments

The authors gratefully acknowledge strategic investment from the Biotechnology & Biological Sciences Research Council (grant reference BBS/E/D/20002173 and tenure-track fellowship support for N.N.L.) and an Academy of Medical Sciences Springboard Award (grant reference SBF007\100134). The funding sources had no role in the design or execution of the experiments described.

We thank Shiranee Sriskandan (Imperial College London), Claire Turner (University of Sheffield) and Alex McCarthy (Imperial College London) for providing GAS isolates. We thank Michael Wessels (Boston Children’s Hospital/Harvard Medical School) and Helge Dorfmueller (University of Dundee) for providing plasmids.

## Supplemental Tables

**Table S1.** Strains used in this study.

| Strain | Description | Reference/source |
| --- | --- | --- |
| *emm1*/TRD<sub>AG</sub> | Identical to H584 | 18 |
| *emm1*_CovS<sub>1-123</sub> | Identical to H598 | 18 |
| *emm1*_5448 | Identical to 5448 | 19 |
| *emm1*<sub>global</sub> | Identical to H1489 | 5 |
| *emm1*<sub>UK</sub> | Identical to H1490 | 5 |
| *emm4*/TRD<sub>AF</sub> |  | Kind Gift, CE Turner |
| *emm5*/TRD<sub>FA</sub> | Identical to Manfredo | 21 |
| *emm12*/TRD<sub>AG</sub> | Identical to H873 | Kind gift, S Sriskandan |
| *emm18*/TRD<sub>DA</sub> | Identical to H566 | 22 |
| *emm49*/TRD<sub>CF</sub> | Identical to NZ131 | 23 |
| *emm89*/TRD<sub>BG</sub> | Identical to H293 | 14 |
| *emm89*/TRD<sub>AG</sub> | 5'TRD<sub>BG</sub> to 5'TRD<sub>AG</sub> | This study |

**Table S2.**
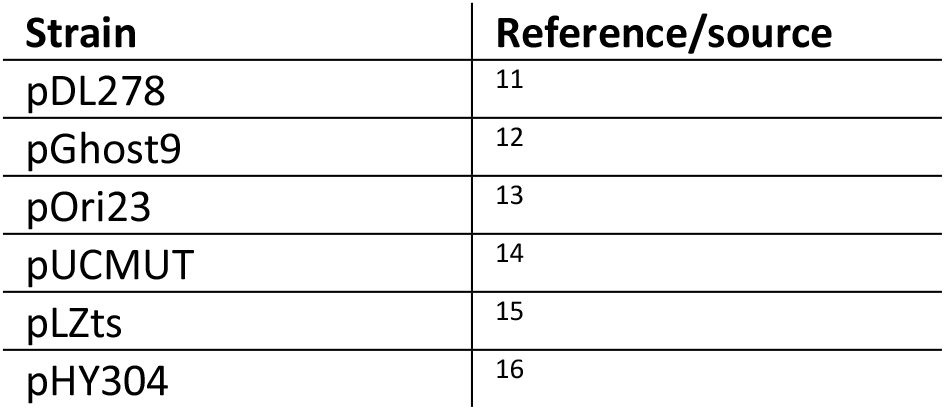
Plasmids used in this study.

| Strain | Reference/source |
| --- | --- |
| pDL278 | 11 |
| pGhost9 | 12 |
| pOri23 | 13 |
| pUCMUT | 14 |
| pLZts | 15 |
| pHY304 | 16 |

**Table S3.**
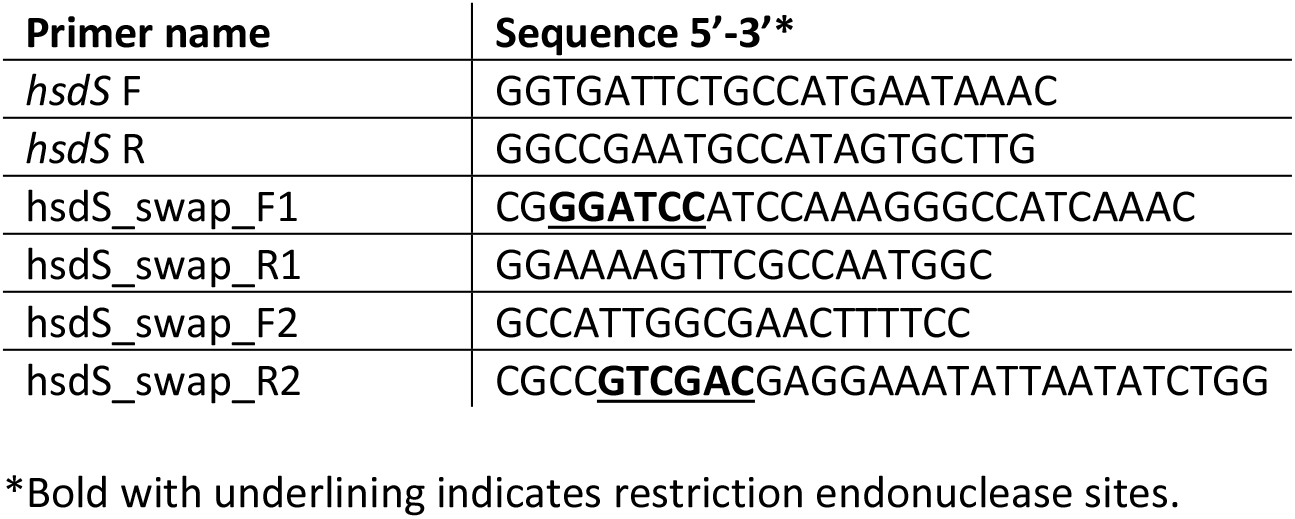
Primers used in this study.

## Supplemental Methods

### Bacterial strains and growth conditions

GAS clinical strains (Table S1) were cultured on trypticase soy agar supplemented with 5% defibrinated sheep blood (R&D) or Todd-Hewitt (TH) (Oxoid) agar or in TH broth at 37°C. *E. coli* strain DH5α (New England Biolabs) was used for storage and passaging of all plasmids and cloning. *E. coli* was cultured on LB agar (Formedium) or broth (Sigma) at 37°C with shaking at 180 rpm. Growth media were supplemented with antibiotics where appropriate at the following concentrations: for *E. coli,* spectinomycin (Merck) at 50 μg/ml, erythromycin (Sigma) at 250 μg/ml; for GAS, spectinomycin at 50 μg/ml and erythromycin at 1 μg/ml.

### Sanger sequencing of the *hsdS* gene

Genomic DNA was extracted from GAS cultures grown to late logarithmic growth phase (OD600 0.7-0.9) as described previously^24^. PCR was carried out with primers *hsdS* F and *hsdS* R (Table S3) with Phusion Flash High-Fidelity PCR Master Mix (Thermo) using a MyCycler (Bio-Rad) thermal cycler. Sanger sequencing was performed in order to determine the sequence of the *hsdS* gene.

### Transformation protocol

#### Preparation of competent cells

GAS were cultured to OD600 0.2 in 40 ml THB and pellets washed 5 times in 1 ml ice-cold 0.5 M sucrose (Sigma) (16000 xg, 1 minute, 4°C). Pellets were re-suspended in a final volume of 100 μl ice-cold 0.5 M sucrose and 50 μl aliquots were used immediately for electrotoporation^7^.

#### Electroporation

5 μl plasmid DNA (at 100 ng/μl) was mixed with 50 μl competent cells and stored on ice. DNA was transformed by electroporation (MicroPulser Electroporator, Biorad) with the following settings: 200 Ω, 1.7 kV, 50 μF, 0.1 cm cuvette (Thermo). Cells were recovered in 1 ml THB, cultured for 1 hour at 37°C for pDL278 or 2 hours at 30 for pJRS233 and pGhost9, and plated on selective media^7^. Where relevant 1 μl (2.5 μg) Ocr (TypeOne™, Lucigen) was added to electroporation reactions prior to pulsing.

### Purification of GAS self-methylated plasmid

Plasmid DNA was purified from successfully transformed GAS strains using a modified QIAprep Spin Miniprep (Qiagen) purification protocol. Briefly, GAS were cultured overnight in 50 ml THB and pellets resuspended in 1 ml QIAprep buffer P1 supplemented with mutanolysin (100 units/ml, Sigma) and lysozyme (1 mg/ml), and incubated for 30 minutes at 37°C. Lysates were divided into 4x 250 μl aliquots and mixed with buffers P2 and N3 as per manufacturers guidelines. Following a 10-minute centrifugation step, supernatants were concentrated 2-fold and then purified twice over sequential QIAprep columns. Plasmid DNA was eluted in 50 μl nuclease-free water/column.

### Generation of TRD_AG_/TRD_BG_ swap strain

Using the temperature-sensitive *E. coli-GAS* shuttle vector pJRS233^20^, plasmid pJRS_hsdS_M89/M1_swap was generated to facilitate creation of an isogenic *emm*89/TRD_BG_ strain expressing the TRD_AG_ *hsdS* allele, where the 5’ TRD_A_ was swapped with TRD_B_. Primers hsdS_swap_F1 and _R1 (Table S3) were used to amplify the 5’ TRD_A_ sequence from genomic DNA purified from *emm*1 GAS. A region of DNA downstream of the TRD was necessary to facilitate homologous recombination within the *emm*89 genome. Due to a single SNP in the 3’ homologous region, this second amplicon was amplified from *emm*89 genomic DNA using primers hsdS_swap_F1 and _R1 (Table S3) and spliced to 5’ TRD_A_ by overlap extension (SOEing) PCR, using overlapping the amplicons as target DNA. This was performed using primers hsdS_swap_F1 and _R2 (Table S3), incorporating the restriction sites BamHI and SalI into the resulting PCR product to facilitate cloning into the vector pJRS233. The resulting shuttle vector pJRS_hsdS_M89/M1_swap was transformed into GAS-M1 strain 854 by electroporation and 5’TRD_A_ was exchanged with the chromosomal copy of 5’TRD_B_ by allelic-exchange mutagenesis (43). Sanger sequencing was performed to confirm introduction of TRD_AG_ into the chromosomal TRD_BG_ gene.

